# Examining the Role of the Surfactant Family Member SFTA3 in Interneuron Specification

**DOI:** 10.1101/330621

**Authors:** Christopher Y. Chen, Nickesha C. Anderson, Sandy Becker, Martin Schicht, Christopher Stoddard, Lars Bräuer, Friedrich Paulsen, Laura Grabel

## Abstract

The transcription factor *NKX2.1*, expressed at high levels in the medial ganglionic eminence (MGE), is a master regulator of cortical interneuron progenitor development. To identify gene candidates with expression profiles similar to *NKX2.1*, previous transcriptome analysis of human embryonic stem cell (hESC)-derived MGE-like progenitors revealed *SFTA3* as the strongest candidate. Quantitative real-time PCR analysis of hESC-derived NKX2.1-positive progenitors and transcriptome data available from the Allen Institute for Brain Science revealed comparable expression patterns for *NKX2.1* and *SFTA3* during interneuron differentiation *in vitro* and demonstrated high *SFTA3* expression in the human MGE. Although *SFTA3* has been well studied in the lung, the possible role of this surfactant protein in the MGE during embryonic development remains unexamined. To determine if *SFTA3* plays a role in MGE specification, *SFTA3*-/- and *NKX2*.1 -/-hESC lines were generated using custom designed CRISPRs. We show that *NKX2.1* KOs have a significantly diminished capacity to differentiate into MGE interneuron subtypes. *SFTA3* KOs also demonstrated a somewhat reduced ability to differentiate down the MGE-like lineage, although not as severe relative to *NKX2.1* deficiency. These results suggest *NKX2.1* and *SFTA3* are co-regulated genes, and that deletion of *SFTA3* does not lead to a major change in the specification of MGE derivatives.

## Introduction

During early embryonic development of the mammalian telencephalon, the transcription factor *NKX2.1* is highly expressed in the medial ganglionic eminence (MGE), a subpallidal structure of the ventral forebrain (1-3). The MGE and caudal ganglionic eminence (CGE) are transient embryonic structures that are the primary source of GABAergic inhibitory progenitors, which migrate tangentially to target sites in the cortex. These progenitors then differentiate into a number of diverse inhibitory interneuron subtypes that modulate the activity of excitatory projection neurons in the cerebral cortex (3-8). Expression of the homeobox protein NKX2.1 is a requirement for specification of the MGE and its derivatives. *NKX2.1* deficient mice display gross malformations of the ganglionic eminences and a complete loss of specific MGE-derived subtypes such as parvalbumin (PV) and somatostatin (SST) expressing interneurons (Butt et al., 2008; Du et al., 2008; Ohkubo et al., 2002). The experimental downregulation of *NKX2.1* in the ventral subpallium results in a conversion of MGE to CGE fates. In *NKX2.1* conditional loss-of-function studies in mice, an increase in the generation of vasoactive intestinal polypeptide (VIP) and calretinin (CR)-expressing interneurons derived from the CGE is produced at the expense of MGE subtypes (9-11). These results indicate NKX2.1 is a master regulator that establishes the MGE and promotes specification of interneuron subtypes.

To identify gene candidates with expression profiles similar to *NKX2.1* that could also help specify the MGE lineage, we utilized previously published data from our laboratory comparing the RNAseq-based transcriptome of FACS isolated human embryonic stem cell (hESC) -derived NKX2.1-positive progenitors to NKX2.1-negative cells. This analysis showed that the profile of surfactant associated 3 (*SFTA3*) expression closely followed *NKX2.1*, suggesting it was a novel MGE marker that could be involved in interneuron specification (12).

*SFTA3*, which encodes surfactant protein H, is part of the multifunctional surfactant gene family, implicated in immune host defense and regulation of alveolar surface tension during normal respiratory mechanics of the lung. To date, there are four surfactant proteins, A, B, C, and D, that have been extensively characterized in the lung. Surfactant proteins A and D are associated with the collectin gene family and are implicated in immunoregulatory host defense. These proteins contain a lectin domain to allow surfactant binding of viruses, fungi, and bacteria, facilitating opsonization for phagocytic digestion and removal (13, 14). In contrast, SP-B and SP-C are hydrophobic proteins required for stabilization of the air-liquid interface at the lung surface to prevent collapse of the alveoli (15, 16). Recently, SFTA3 was identified by bioinformatics as a novel secretory surfactant protein expressed in the human lung (17). Interestingly, SFTA3 shares very little sequence or structural similarity when compared to other surfactants or proteins in general. The BLAST search tool algorithm in conjunction with the Uniprot protein database revealed SP-H homologs in primate species only, including humans. No matches were found when additional BLAST searches were performed comparing the SP-H sequence to 3D biochemical structures listed in the Protein Data Bank (17).

Numerous sequence-based prediction tools used to identify post-translational modification (PTM) sites in the SP-H sequence suggested a high probability of palmitoylation, glycosylation, and phosphorylation. These data suggest that SFTA3 is an amphiphilic protein that can acquire both hydrophobic and hydrophilic properties (17). Western blot analysis using an anti-SP-H antibody in the A549 alveolar lung cell line detected a distinct band for the expressed protein at ∼13kDa (17). Similar to the other proteins in the surfactant family, studies demonstrate that SFTA3 has a role in innate immune response localized to the lipid plasma membrane surface (17, 18). Computer simulations investigating the binding affinity of SP-H with dipalmitoylphosphatidylcholine (DPPC), the most prevalent lipid in pulmonary surfactant, illustrated protein stability for SP-H at the lipid surface (19). These data suggest PTMs are responsible for the amphiphilic properties of *SFTA3* in specifying either surface regulatory or immune defense function.

*SFTA3* is a single copy gene immediately adjacent to *NKX2.1* on chromosome 14q13.3. This location, combined with their correlated temporal and spatial patterns of expression suggest that *NKX2.1* and *SFTA3* may regulate common developmental pathways. For example, *NKX2.1* is also expressed during early development of the lungs and is implicated in promoting the production of surfactants in alveolar cells. The production of surfactant is perturbed upon disruption of the *NKX2.1* gene (20). Moreover, patients with mutations in *SFTA3* display a variety of aberrant symptomology including choreoathetosis, hypothyroidism, and neonatal respiratory disease (21). Approximately 50% of patients with mutations in *NKX2.1* develop the same clinical phenotypes of motor ataxia and respiratory distress, all part of a larger connected network of disorders known as “brain-lung-thyroid syndrome” (OMIM 610978) (22). The mouse orthologue of *SFTA3* is NKX2.1-associated noncoding intergenic RNA (NANCI). Recent studies demonstrate a regulatory role for NANCI in *NKX2.1* expression in the mouse lung (23, 24). Intriguingly, whereas the human *SFTA3* gene contains an apparent open reading frame that is translated, mouse NANCI encodes a long non-coding RNA with no apparent open reading frames. The potential interaction between *NKX2.1* and *SFTA3* is largely unexamined, and a function for *SFTA3* outside of the lung has not been established.

We now show using quantitative PCR (qRT-PCR) analysis upregulation in *SFTA3* gene expression during differentiation of hESC-derived progenitors to an MGE-like fate. The BrainSpan Atlas, an open source database using RNA-sequencing to profile cortical and subcortical structures at various time points during early embryonic development, indicates *SFTA3* expression is selectively upregulated in the MGE at 8-9 weeks gestation. We generated *SFTA3* and *NKX2.*1 knock-out (KO) hESC lines using CRISPR-Cas9 genome editing in order to examine two fundamental questions. First, to determine if *NKX2.1* and *SFTA3* genes are co-regulated such that the deletion of one gene will modify the expression pattern of the other. Second, to determine if *SFTA3* serves a functional role in the specification of MGE GABAergic progenitors and their differentiation into mature inhibitory interneuron subtypes. An *NKX2.1* KO cell line, expected to be deficient in specifying MGE-derived interneuron subtypes, served as a control to determine *SFTA3* function in specifying MGE lineage identity. The deletion of *NKX2.1* resulted in a dramatic reduction in *SFTA3* expression and a significant decline in the number of cells expressing SP-H. In contrast, the loss of *SFTA3* led to no decline in *NKX2.1* message and a slight decrease in the number of cells expressing NKX2.1 protein. Additionally, the absence of *NKX2.1* resulted in the virtual elimination of MGE-like gene expression with a concomitant increase in expression of non-MGE cell fates including dorsal forebrain and CGE phenotypes. Mutations to *SFTA3* resulted only in a moderate decrease in MGE-associated gene expression with no concomitant increase to non-MGE derivatives.

## Materials and Methods

### Generation of KO cell lines

A dual sgRNA-directed gene knockout approach using CRISPR-Cas9 was used to remove a portion of the *SFTA3* gene. One guide sequence was designed to cut upstream of exon 2 and a second guide sequence was designed to cut downstream of exon 2. Deletion of exon 2 resulted in an early termination and a severely truncated protein. For deletion of *NKX2.1*, two sgRNA-directed CAS9 nuclease were targeted to flank the entire gene to remove all isoforms of NKX2.1, completely removing all exons from the genome. Guide RNA sequences were cloned into addgene plasmid #62988. The dual sgRNA vector was electroporated into H9 hESC cells using a Gene Pulser X (250 V, 500 uF). Cells were plated and after 24 hours, puromycin was supplemented into the medium at 1 ug/ml. Puromycin was added for a total of 48 hours to select for the transient expression of the dual sgRNA vectors. After 14 days, clones were isolated, expanded and tested by PCR. Clones were screened for deletion, inversion, and zygosity by PCR. Double KO clones were further expanded and sequenced across the junction to confirm genomic deletion. The same procedures described above were performed to derive control lines with the exception that no gRNA sequence was used during vector electroporation.

### Culture of ESCs and MGE-like cells

hESC *SFTA3* and *NKX2.1* control and KO cell lines were maintained and passaged as previously described (25). Neural differentiation of ESCs was initiated using the ALK2/3 inhibitor LDN-193189 (Stemgent, 100 nM) and progenitors ventralized to an MGE-like identity by a combination of sonic hedgehog (R&D Systems, 125 ng/mL) and its agonist purmorphamine (Calbiochem, 1 µM) as previously described (26).

### ***In vitro* maturation of hESNPs**

The addition of the ROCK inhibitor Y27632 (1 µM, Calbiochem) was supplemented into every medium change of *SFTA3* control and KO progenitors beginning at differentiation day 21 until day of fixation.

### Immunocytochemistry

The fixation, permeabilization, and incubation of primary and secondary antibodies were performed as previously described (Chen et al., 2016). The following antibodies were used: DLX2 (Proteintech, rabbit, 1:50), Olig2 (Proteintech, rabbit, 1:100), DCX (Millipore, guinea pig, 1:500), FOXG1 (abcam, rabbit, 1:100), GABA (Sigma, rabbit, 1:500), MAP2 (Sigma, mouse, 1:1000), NKX2.1 (Chemicon, mouse, 1:250), and Nestin (Millipore, mouse, 1:1000). Hoechst 3342 (Molecular Probes) was used to counterstain all cell nuclei and slides coverslipped with gelvatol. Images used for quantification were taken on a Nikon Eclipse Ti microscope with NIS-Elements software.

### Quantitative Real Time PCR

Quantitative measurements of mRNA expression were performed using the 7300 Real Time PCR System (Applied Biosystems) as previously described (12).

### RNA Sequencing and Bioinformatics Analyses

Characterization of the transcriptomes comparing NKX2.1-positive and NKX2.1-negative populations using shotgun mRNA sequencing (RNA-seq) was performed as previously described (12). RNA-Seq data from the BrainSpan Atlas of the Developing Human Brain (http://brainspan.org) was assessed as previously described (12).

## Results

### Comparative Gene Expression Analysis of SFTA3 and NKX2.1 in MGE-like interneuron progenitors and Fetal Brain tissue

We used RNA-sequencing analysis to compare the transcriptome of NKX2.1-positive and NKX2.1negative neural progenitors to identify genes whose expression was enriched in the NKX2.1-positive MGE-like population. SFTA3 expression was consistently enriched in the NKX2.1-positive population, and to the same extent observed for NKX2.1 (>8 fold gene enrichment) (Chen et al., 2016; Figure 1A). The expression values for *SFTA3* are comparable to *NKX2.1*, with a correlative-coefficient R value >0.98; indicating strong statistical significance in fold change enrichment between these two genes (Supplementary Table 1).

**Figure 1:**
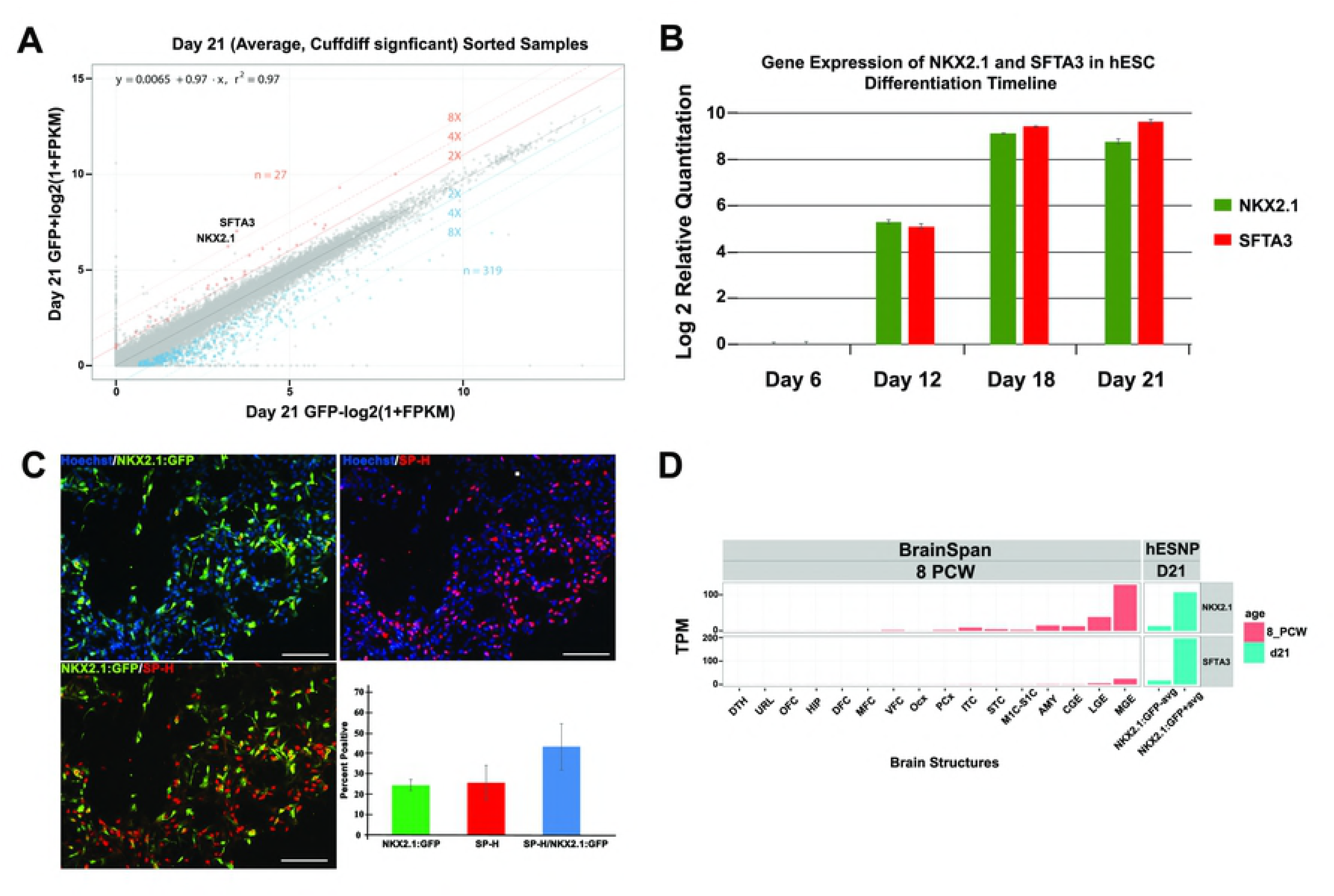
Comparative gene expression analysis of *SFTA3* and *NKX2.1*. A) *SFTA3* and *NKX2.1* pairwise gene expression analysis indicates significant levels of enrichment in the NKX2.1-positve progenitor population in comparison to the NKX2.1-negative cohort. Fold change differences of 2, 4, and 8 between samples are represented by central diagonal lines. Orange and blue points signify enriched and depleted differentially expressed genes (FDR<0.05) comparing NKX2.1-positive to NKX2.1-negative progenitor populations with the total number of genes indicated for each category. B) RT-PCR analysis of *SFTA3* and *NKX2.1* gene expression at specific time points during the hESC differentiation timeline. C) *In vitro* day 26 immunocytochemistry and quantification of neural progenitors for NKX2.1 and SP-H. Data represented as ± SEM. Scale bar = 100 µm. D) RNA-seq comparison of FACS isolated NKX2.1positive and NKX2.1-negative cell populations in comparison to the developmental transcriptome of various structures of the human brain at 8 and 9 pcw. Brain structure legend: DTH, dorsal thalamus; DFC, dorsolateral prefrontal cortex; HIP, hippocampus; OFC, orbital frontal cortex; Ocx, occipital neocortex; MFC, anterior cingulate cortex; PCx, parietal neocortex; URL. upper rhombic lip; VFC, ventrolateral prefrontal cortex; STC, posterior superior temporal cortex; M1C-S1C, primary motor-sensory cortex; ITC, inferolateral temporal cortex; AMY, amygaloid complex; CGE, caudal ganglionic eminence; LGE, lateral ganglionic eminence; MGE, medial ganglionic eminence.

We used RT-PCR analysis to examine expression levels of these two genes at specific time points during the differentiation of hESC-derived interneurons. Both genes were upregulated in a similar time course, consistent with shared regulatory elements (Figure 1B). Recently, an enhancer sequence region that may confer ventral forebrain expression was identified on chromosome 14 near the *NKX2.1* and *SFTA3* genes (Patent Seq ID No. 144). This shared enhancer region may be responsible for upregulating the transcription of both *SFTA3* and *NKX2.1*. Additionally, immunocytochemistry revealed a large percentage of neural progenitors at day 26 of differentiation co-expressed NKX2.1 and SP-H (Figure 1C).

To investigate whether *SFTA3* is expressed during the development of the human fetal brain, we examined transcriptome data from the BrainSpan Atlas. The Atlas indicates enrichment of *SFTA3* in the MGE at 8 post-conception weeks (pcw) (Figure 1D). Furthermore, when comparing the expression levels of *SFTA3* to all brain structures at 8 pcw, the MGE is the only structure to demonstrate a significant elevation in expression of the gene (Figure 1D). These results suggest that *SFTA3* is a novel biomarker specific to the MGE during early ventral forebrain development.

### Day 25 Characterization of SFTA3 and NKX2.1 Knock-out Neural Progenitors

Given the highly correlative spatiotemporal gene expression patterns of *NKX2.1* and *SFTA3* during *in vitro* differentiation of FACS-enriched NKX2.1-postive neural progenitors and in the human MGE at 8 pcw (Figure 1), we determined whether *SFTA3* plays a role in specifying MGE-like neural progenitors, as observed for *NKX2.1*. To investigate SFTA3 gene function, *NKX2.1* and *SFTA3* control and knockout hESCs were generated using CRISPR-Cas9 mediated genome editing. To verify successful KO of each gene, we performed qRT-PCR at day 25 of differentiation, when both genes are expressed at significant levels. Day 25 qRT-PCR analysis of NKX2.1 control and KO progenitors indicated no detectable RNA transcripts of *NKX2.1* in the KO cell line (Supplementary Figure 1A). In addition, the deletion of *NKX2.1* significantly reduced *SFTA3* transcript levels, suggesting *NKX2.1* is needed to maintain control levels of *SFTA3* gene expression (Supplementary Figure 1A). Both *SFTA3* KO cell lines had no detectable RNA transcripts for *SFTA3*. Interestingly, *NKX2.1* RNA expression is not significantly affected in *SFTA3* KO lines (Supplementary Figure 1B; Figure 2B). We also examined levels of protein expression for these two genes. Immunocytochemistry at differentiation day 30, revealed the absence of NKX2.1 and SP-H protein in their respective KO cell lines (Supplementary Figure 1C). Whereas qRT-PCR showed little change in *NKX2.1* expression in *SFTA3* KOs, both *SFTA3* KO lines had lower levels of NKX2.1 protein (Figure 2C, D).

**Figure 2:**
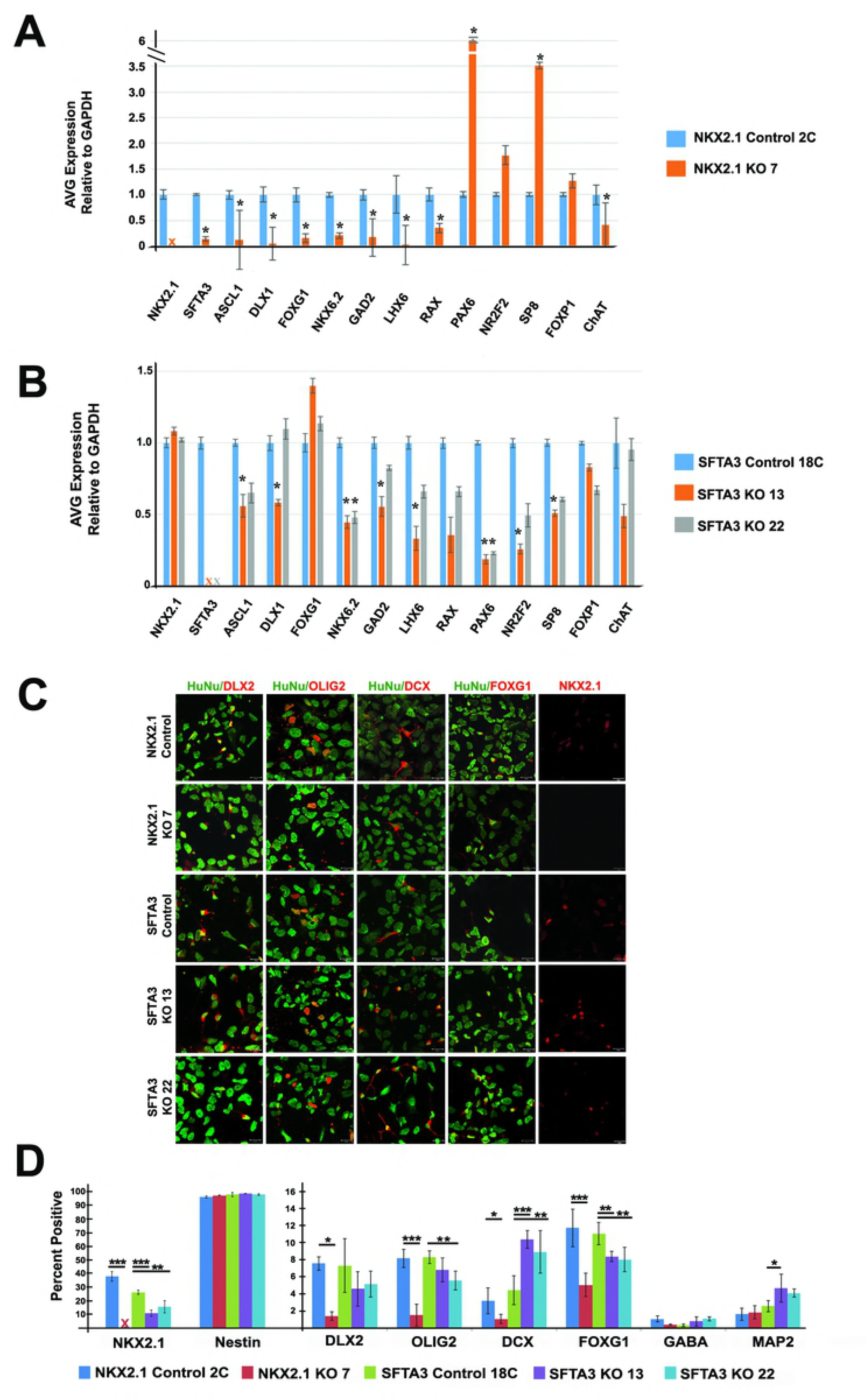
Characterization of day 25 *SFTA3* and *NKX2.1* KO neural progenitors from hESCs. A) RTPCR data comparing gene expression levels between *NKX2.1* control and KO cell progenitors. Data represented as mean ± SEM. * = p<0.05. B) RT-PCR data comparing gene expression levels between SFTA3 control and KO cell progenitors. Data represented as mean ± SEM. * = p<0.05. C) Composite confocal microscopy images that show day 25 immunolabeling of control and knockout hESNPs following differentiation to MGE-like fate. All hESC-derived cells (HuNu, green) and other markers in red are indicated in each panel. Markers label progenitors of the ventral forebrain (NKX2.1, DLX2, Olig2), immature neurons (DCX), neural stem cells (nestin), and telencephalic progenitors (FOXG1). Scale bar = 20 µm. D) Quantification of day 25 immunocytochemistry. Data represented as mean ± SEM. *** = p<0.001, ** = p<0.01, * = p<0.05 (ANOVA).

To investigate *SFTA3* gene function in the specification of MGE cell fate, *NKX2.1* and *SFTA3* control and knockout hESCs were differentiated to MGE-like neural progenitors *in vitro* for 25 days and analyzed for expression of several neural markers using qRT-PCR. Again, no *NKX2.1* was detected in the KO cell line, verifying gene deletion (Figure 2A). Markers of the MGE (*SFTA3, DLX1, NKX6.2, LHX6)*, ventral telencephalon (*ASCL1, RAX*) and prosencephalon (*FOXG1, GAD2,*) displayed significantly decreased levels of expression in the *NKX2.1* KO relative to the control cell line (Figure 2A). Expression of choline acetyltransferase (*ChAT*), an enzyme that synthesizes acetylcholine in cholinergic neurons, also exhibited a significant decrease in the *NKX2.1* KO compared to the control. In addition, expression of markers of dorsal telencephalon and CGE lineage (*Pax6, NR2F2, SP8*) were enriched in the *NKX2.1* KO relative to controls (Figure 2A). There was no significant difference in expression levels of *FOXP1*, a marker of hypothalamic neurons, in the *NKX2.1* KO relative to its control (Figure 2A, B). These data are consistent with *in vivo* studies that demonstrate a re-specification of MGE to non-MGE cell neural fates in conditional *NKX2.1* KO mutant mice (3, 9, 27, 28)

No detectable *SFTA3* mRNA transcripts were observed in the two *SFTA3* KO cell lines, verifying gene mutation (Figure 2B). Though the *SFTA3* KOs showed a general trend towards decreased gene expression of markers of the ventral telencephalon and MGE (*Ascl1, DLX1, NKX6.2, GAD2*, and *LHX6*), the downward trend was not as severe as observed for the *NKX2.1* KOs. Intriguingly, contrary to what was observed in the *NKX2.1* KO cell population, there was no compensatory increase of dorsal forebrain and CGE neural lineage markers (*PAX6, NR2F2, SP8*) in the *SFTA3* KO cells (Figure 2B). These data suggest a discrepancy in phenotype between *NKX2.1* and *SFTA3* KO RNA expression levels, suggesting *SFTA3* is not solely responsible for conferring MGE fate downstream of *NKX2.1*.

We next examined the expression of neural stem cell (NSC), immature neuronal, and ventral progenitor markers in both our control and KO generated cell lines. Greater than 95% of the cells at day 25 in all cell lines expressed the NSC marker nestin, indicating that KO of *NKX2.1* or *SFTA3* did not interfere with neural differentiation (Figure. 2D). While 7.57%, 8.17%, and 11.74% of the *NKX2.1* control cells expressed DLX2, Olig2, and FOXG1 respectively, only 1.40%, 1.51%, and 5.04% of cells expressed these markers in the *NKX2.1* KO cell population, indicating a significant reduction in interneuron progenitor differentiation (Figure 2C, D). *SFTA3* KO progenitors did not display the same decline in comparison to the control cells. Intriguingly, there was a marginal but significant decrease in cells expressing NKX2.1, OLIG2, and FOXG1 protein expression for both *SFTA3* KO lines in comparison to the control (Figure 2C, D). Consistent with the high percentage of cells expressing immature NSC markers at day 25, there was no significant expression of the inhibitory neurotransmitter GABA at this time point (<2%; Figure 2D). Surprisingly, the percentage of cells expressing DCX, a postmitotic neuronal migration marker and MAP2, a mature neuronal marker, was somewhat higher in the *SFTA3* KO lines (8-11%; 4-5%, respectively) compared to its control (4.4%; 2.6%, respectively; Figure 2D). This suggests *SFTA3* may play a role in preventing premature differentiation of neural progenitors and keeping them at the NSC stage. As observed with RNA data, declines in levels of cells expressing MGE and forebrain protein markers were not as dramatic for the *SFTA3* KOs as the *NKX2.1* KOs relative to controls. Overall, the mutation of *SFTA3* alone was not sufficient to diminish expression of MGE and subpallidal markers to *NKX2.1* KO levels, as *SFTA3* KOs showed only a moderate reduction in NKX2.1 expression. Taken together, these results suggest that *SFTA3* does not have a central role in specifying MGE-like progenitor cell lineage as established for *NKX2.1*.

### Day 45 Characterization of SFTA3 and NKX2.1 Knock-Out Neural Cell Populations

To evaluate whether the absence of *SFTA3* affects the differentiation of hESC-derived GABAergic progenitors into inhibitory interneurons, neural progenitor cells were cultured for an additional 20 days and analyzed at day 45 *in vitro*. As hESC-derived interneurons demonstrate a protracted developmental timeline for maturation, maintaining long-term cultures for immunocytochemical analysis was difficult, with some cell lines demonstrating decreased viability at time of analysis. The loss of *NKX2.1* or *SFTA3* expression did not significantly affect maturation of the neural progenitors, as there was comparable expression of both DCX and MAP2 in all KO lines compared to controls (Figure. 3 A, B). Day 45 neurons generated from the NKX2.1 KO line exhibited minimal GABA expression, (< 1%) relative to control cells at this time point (> 38%), consistent with a role for NKX2.1 in conferring GABAergic inhibitory identity. Immunocytochemistry-based quantification of *SFTA3* mutants indicated ∼10% GABA expression in both *SFTA3* knockout lines compared to ∼18% expression in the control, demonstrating that the deletion of *SFTA3* did not eliminate neural progenitor differentiation down the GABAergic lineage (Figure 3A, B).

**Figure 3:**
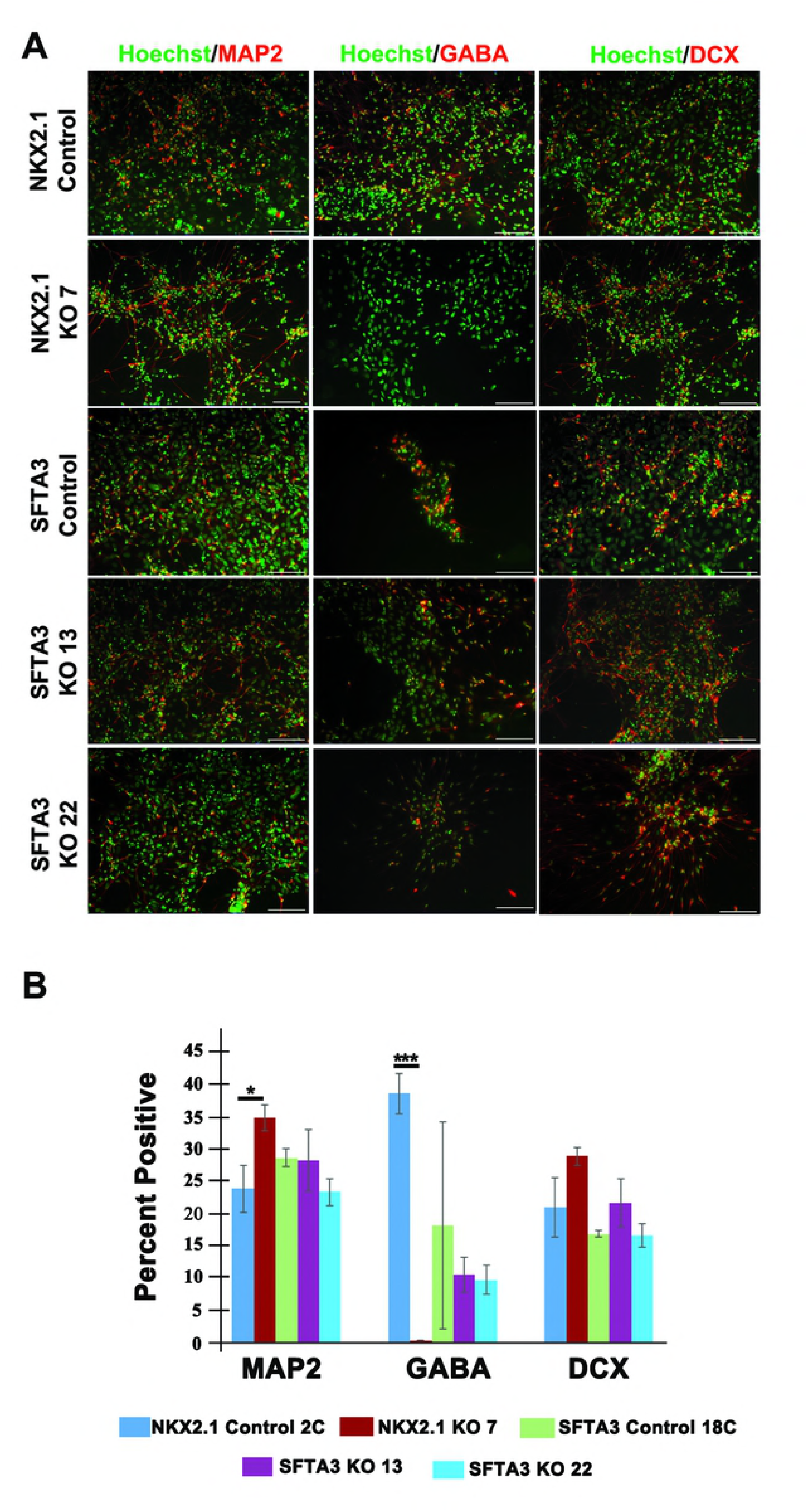
Characterization of mature hESC-derived neurons from day 45 *SFTA3* and *NKX2.1* KO lines. Composite micrographs representing day 45 immunocytochemistry of mature inhibitory interneurons differentiated from control and KO hESNPs. Immunolabeling for markers of immature neuronal precursor cells (DCX), mature neurons (MAP2) and GABAergic neurons (GABA). Scale bar =100 µm. Day 45 immunocytochemistry quantification analysis of GABA, DCX, and MAP2 expression. Data represented as ± SEM. *** = p<0.001, ** = p<0.01, * = p<0.05 (ANOVA).

## Discussion

The transcription factor *NKX2.1* is a master regulator of MGE-specific lineages, promoting the expression of genes involved in specifying cortical inhibitory interneuron progenitors and helping to maintain proper ratios of mature interneuron subtypes in the adult brain (3, 29, 30). This report highlights the identification of a surfactant gene that is expressed in NKX2.1-positive cells and may support inhibitory interneuron differentiation, though not as robustly as *NKX2.1*. Based upon transcriptome and qRT-PCR analysis, we identify *SFTA3* as a novel MGE marker (Figure 1) and examine *SFTA3*’s role in interneuron differentiation by generating *NKX2.1* and *SFTA3* KO hESC lines and determining their ability to produce MGE-subtypes and derivatives. We show that MGE-lineage specific markers and interneuron subtypes are only marginally decreased when *SFTA3* is mutated, whereas the *NKX2.1* KO showed significant depletion in both GABAergic neural progenitor and cortical inhibitory interneurons. By day 25 of differentiation, *NKX2.1* KO progenitors showed significant levels of enrichment in expression of dorsal and CGE-associated genes at the expense of MGE marker expression (Figure 2A). Surprisingly, at day 45, despite our observation that non-MGE like interneuron progenitors were present at day 25, no cells expressing GABA were detected in the *NKX2.1* KO. This suggests the non-MGE alternative fated cells specified at day 25 could not mature under our *in vitro* conditions. (Figure 3A, B; Butt et al., 2008; Sussel et al., 1999).

Analysis of RNA expression levels for *ASCL1*, a proneural gene associated with promoting cell cycle exit and neuronal differentiation was reduced in both *SFTA3* KO lines in comparison to the control. Additionally, immunocytochemistry data interestingly showed a significant increase in the percentage of cells that expressed DCX and MAP2 in the SFTA3 KOs relative to the control at day 25 of differentiation. These results highlight a possible function of *SFTA3* as a cell-cycle regulator of MGE-like progenitors during neurogenesis.

The orthologue of the human *SFTA3* gene in rodents is NANCI, which appears to encode a long noncoding RNA (lncRNA). A comparison of coding sequences using nucleotide BLAST algorithms was performed in order to understand why human SFTA3 is a protein encoding gene and mouse NANCI a lncRNA (data not shown). The NANCI coding sequence lacks long open reading frames due to the presence of numerous translational stop codons interspersed throughout the sequence. In contrast, the human SFTA3 coding sequence contains long uninterrupted open reading frames that do not contain stop codons, consistent with protein coding regions. Recently, Herriges and colleagues observed nearly identical expression patterns during development for both NANCI and *NKX2.1* in the mouse lung epithelium and forebrain (23). Heterozygous NANCI mouse mutants had decreased levels of *NKX2.1* expression but did not display significant morphological defects in the lungs. Furthermore, heterozygous NKX2.1 mutants showed a compensatory increase in NANCI expression, resulting in an upregulation of *NKX2.1* expression back to wild-type levels (24). These data suggest that NANCI may function as a regulator of *NKX2.1* expression. We observed similar tissue specific *NKX2.*1 and *SFTA3* gene expression patterns in *in vitro* derived MGE-like progenitor cell population and in transcriptome data of human 8 pcw MGE fetal tissue. Despite similarities in the tissue specificity of expression of *SFTA3* and NANCI, our data do not support a role for *SFTA3* in regulating *NKX2.1* expression. Further investigation to verify the presence and potential function of an *SFTA3* long noncoding RNA in human cells is needed.

In summary, our data demonstrate that the expression of *SFTA3* and *NKX2.1* are coordinated during differentiation of hESCs *in vitro* to an MGE-like fate. Deletion of *SFTA3* only marginally affects the expression of *NKX2.1*, and does not lead to any major changes in the expression profiles of MGE markers and interneuron derivatives. Further research is needed to fully assess the functional differences between *SFTA3*/NANCI and *NKX2.1* in rodents and humans.

## Acknowledgements

This work was funded by grant 13-SCC-WES-01 from the Connecticut Regenerative Medicine Research Fund to L. Grabel.

## Supplementary Figure Legends

**Supplemental Figure 1**. Verification of *NKX2.1* and *SFTA3* gene knockout in hESC lines. A) qRT-PCR data comparing *NKX2.1* and *SFTA3* expression levels between day 25 *NKX2.1* control and KO cell progenitors. Data represented as mean ± SEM. * = p<0.05. B) qRT-PCR data comparing *NKX2.1* and *SFTA3* gene expression between day 25 *SFTA3* control and KO cell progenitors. Data represented as mean ± SEM. * = p<0.05. C) Day 25 immunocytochemistry analysis of *NKX2.1* and *SFTA3* knockout and control hESNPs. Scale bar =20 µm

**Supplementary Table 1**. *SFTA3* and *NKX2.1* transcriptome data of correlation and fold change measurements.

